# A reversible KO model reveals therapeutic potentials of defective Tregs

**DOI:** 10.1101/770347

**Authors:** Yongqin Li, Tian Chi

## Abstract

Tregs must be activated to suppress immune responses, but the transcriptional program controlling Treg activation remains incompletely understood. We previously found that Treg-specific deletion of the chromatin remodeling factor *Brg1* impairs Treg activation and causes fatal autoimmunity in mice. Here, using a method that allows gene KO to be reversed in a Tamoxifen-dependent manner, we addressed whether reinstating *Brg1* expression in the defective Tregs in the sick mice could restore Treg function, and if so, whether such Tregs could stop and resolve the fatal inflammation. We found that reexpressing *Brg1* unexpectedly converted the defective Tregs into highly potent “SuperTregs”, which effectively rescued the dying mice. Remarkably, *Brg1* reexpression in as little as 8% of the Tregs sufficed for the rescue in some cases. *Brg1*-deleted Tregs in the inflamed mice experienced excessive cytokine stimulation, became hyperactivated upon *Brg1* reexpression and then deactivated as the inflammation subsided, suggesting that BRG1 acted in conjunction with inflammation to induce and maintain the SuperTreg phenotype. These data illustrate the power of reversible KO models in uncovering gene functions, and suggest a novel therapeutic strategy for IPEX(-related) disorders that exploits the defective Tregs and the inflammatory environment preexisting within the patients.

## INTRODUCTION

Tregs are potent suppressors of immune responses^1,2^, and defects in Treg development and/or function can underlie devastating autoimmune disorders^3–5^. The majority of Tregs under physiological conditions are naïve, with little overt suppressor activity. Upon antigen and cytokine stimulation, naïve Tregs become activated and differentiated into effector cells expressing various cell surface and soluble molecules that mediate suppressor function^1,6–9^. It is therefore of great interest to characterize the mechanisms controlling Treg activation and effector function.

*Brg1* is the catalytic subunit of the chromatin remodeling BAF (mSwi/snf) complex ^10^, which plays diverse roles in the immune system ^11–16^. We have identified *Brg1* as a crucial regulator of Treg activation^17^. Specifically, *Brg1* deletion in Tregs impairs Treg activation, concomitant with the onset of inflammation. Remarkably, as the inflammation progresses, Tregs become increasingly activated, but the activation levels are unable to catch up with the severity of inflammation, which fails to stop the progression of the disease, leading ultimately to the death of the KO mice. These data indicate that *BRG1* acts to sensitize naïve Tregs to inflammatory cues, thus allowing them to promptly and effectively suppress autoimmunity^17^.

Our study described above is focused on the role of *Brg1* in naïve Tregs in the healthy mice. To extend this line of investigation, we sought to determine whether in the *Brg1* KO mice that have developed severe inflammation, reinstating *Brg1* expression in the partially activated, *Brg1*-deleted Tregs could restore Treg function and even rescue the dying mice. There is no reason *a priori* to assume positive answers to these questions. To regulate target genes, *Brg1* must act in conjunction with other transcription regulators including sequence-specific transcription activators and histone modifying enzymes. These other regulators provide the informational context for *Brg1* function, which can dictate the outcome of *Brg1* expression. As this context might differ in naïve vs. (partially) activated Tregs, it is difficult to infer, based on the role of *Brg1* in naïve Tregs, the outcome of *Brg1* reexpression in the partially activated, *Brg1*-deleted Tregs. Even if *Brg1*-reexpression can restore Treg function, it is unclear whether this is sufficient to resolve the severe inflammation and rescue the mice, given that the inflammation may have become overwhelming and/or tissue damages irreversible by the time of *Brg1* reexpression. These considerations are not only important for understanding *Brg1* function, but also have therapeutic implications for human autoimmune diseases resulting from Treg defects (see Discussion).

We have addressed these issues using LOFT, a reversible gene targeting strategy we previously developed^18^. The results reveal dramatic therapeutic effects of Brg reexpression on the sick mice, which is of both biological and clinical interest.

## RESULT

### The LOFT strategy for *Brg1* reversible KO (rKO)

Treg-specific *Brg1* deletion followed by conditional restoration of *Brg1* expression was achieved with the LOFT method ^18^ that requires a pair of alleles of the target gene (*Brg1* in the current study): a floxed allele (*Brg1*^*F*^) and a reversibly trapped allele that is a null by default but can be conditionally converted to a wild-type (WT) allele. The latter allele is designated Δ*R*, where R denotes ‘reversible’ (Figure 1A, top left). The key component of the ΔR allele is a gene-trap cassette consisting of the neomycin phosphotransferase (Neo) and Ires-GFP. This cassette was inserted into intron #9 (Fig. 1B), thus capturing the upstream exon #8 (E8) to produce a fusion protein between the N-terminal 531 aa of *BRG1* protein and the neomycin phosphotransferase, the former moiety being inactive, and the latter serving as the selection marker for successfully targeted embryonic stem (ES) cells. In addition, GFP was co-expressed with the fusion protein, which reported the status of Δ*R* allele. The gene-trap cassette was flanked by FLP recombination target (FRT) sites, allowing for conditional cassette excision in the presence of the FLP recombinase. The removal of the gene-trap cassette restores the expression of full-length *BRG1*, concomitant with the loss of GFP expression. Thus, in *Brg1*^*F/ΔR*^ mice that also expressed Cre in Tregs (from the *FoxP3*^*YFP-Cre*^ *allele) and FlpoER* (from the ubiquitous *CAG* promoter inserted into *R26* locus), *Brg1* expression is constitutively eliminated in Tregs but reinstated upon Tamoxifen (TAM) administration, the latter event reported by elimination of GFP fluorescence (Figure 1A, middle and bottom).

**Fig. 1.**
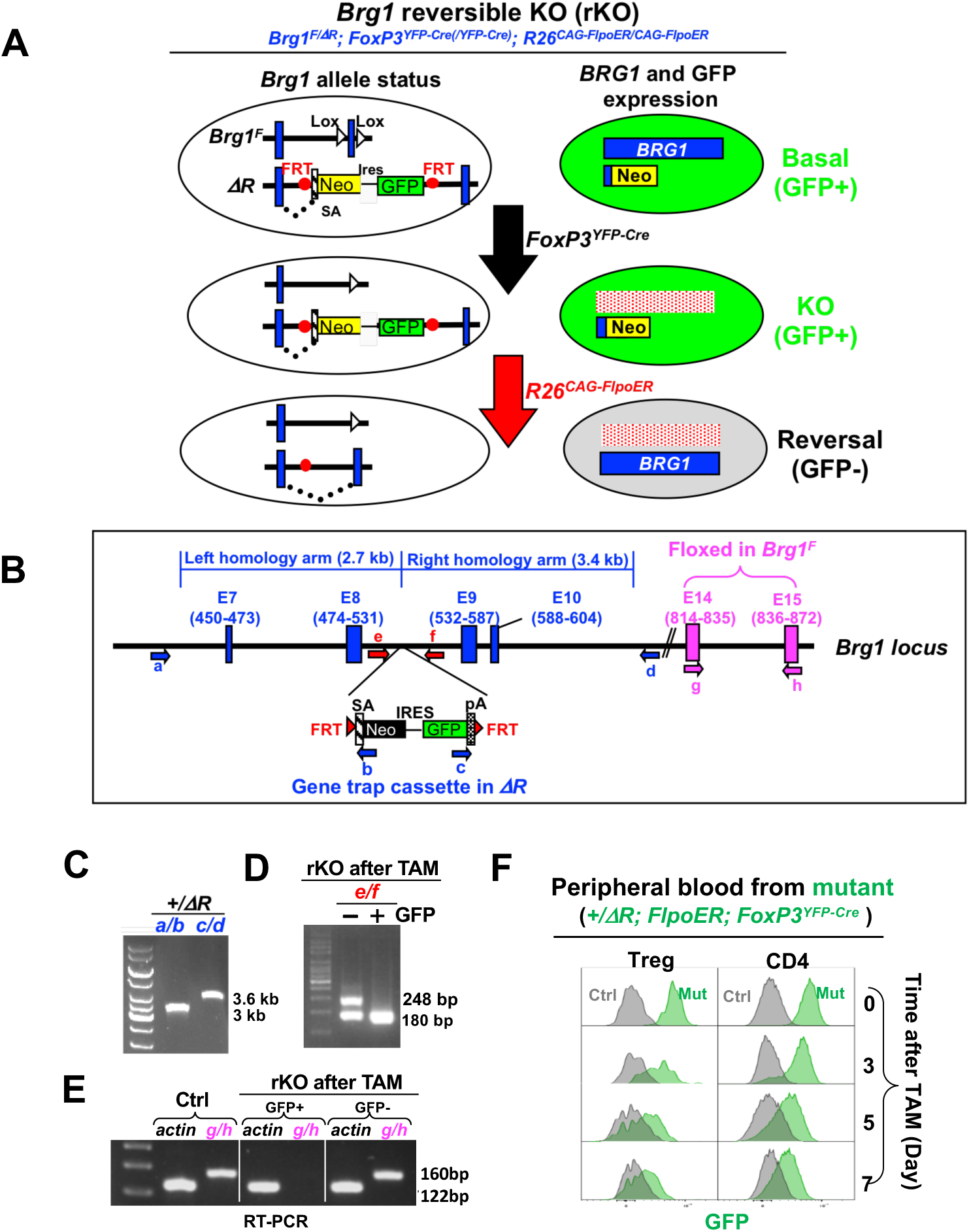
Creation of *Brg1* reversible KO (rKO) model using the LOFT method. **(A)** Strategy for GFP-labeled, Treg-specific reversible *Brg1* knockout. This method requires a conventional *Brg1* floxed allele (*Brg1*^*F*^) paired with a multi-functional reversible KO (ΔR) allele (top left), and sequential action of Cre and Flpo recombinases (middle and bottom). Depicted are the status of the *Brg1* alleles (left) and the corresponding *BRG1* protein expression patterns (right). Note that Cre was expressed from the endogenous FoxP3 locus located on the X chromosome subject to random inactivation, and so the rKO mice carried either one or two *FoxP3*^*YFP-Cre*^ alleles depending on the sex. SA, splicing acceptor; Neo, neomycin resistance gene; FRT, Flippase recognition target (red dot). (B) The *Brg1* alleles. The gene trap cassette in Δ*R* is inserted after E8 in the *Brg1* locus, and the floxed exons in *Brg1*^*F*^ highlighted in pink. Depicted also are the homology arms used to make the targeting construct for generating Δ*R*, and the PCR primers for genotyping. (C-E) Characterization of mouse samples. Tail from a *Brg1*^*+/ΔR*^ mouse was subjected to PCR analysis using primer pair a/b and c/d (depicted in Fig. 1B) to verify successful targeting (C); GFP^+^ and GFP^-^ Tregs isolated from TAM-treated rKO mice were analyzed by PCR and RT-PCR to detect the excision of the gene trap cassette (D) and restoration of *Brg1* expression (E), respectively. The control mouse in (E) has the same genotype as rKO except that it lacked *FoxP3*^*YFP-Cre R*^. (F) Kinetics of GFP loss following TAM administration. TAM (full dose) was given via oral gavage and GFP expression in Tregs and conventional CD4 cells in tail blood monitored by FACS. The control mouse did not carry *Brg1*^Δ*R*^

### Characterization of the *ΔR* allele

We inserted the gene trap cassette into the ES cells using the traditional gene targeting method (Fig. 1B) to generate *Brg1*^*+/ΔR*^; *R26*^*CAG-FlpoER*^ mice. PCR analysis confirmed that the mice carried Δ*R* (Fig. 1C). Following oral gavage of a full dose of TAM (500 ug/g, once daily for two consecutive days, termed the “full dose” regimen hereafter), GFP signal in the Tregs in the peripheral blood decayed gradually, disappearing almost completely on Day 7 after the gavage (Fig. 1F, left), the kinetics being comparable to that in the conventional CD4 cells (Fig. 1F, right). Finally, we bred the rKO mice by introducing *Brg1*^*F*^and *FoxP3*^*YFP-Cre*^ into the *Brg1*^*+/ΔR*^; *R26*^*CAG-FlpoER*^ mice. As *FoxP3*^*YFP-Cre*^ is located on the X chromosome randomly inactivated in females, the genotypes of rKO mice are gender-specific, being *Brg1*^*F/ΔR*^; *FoxP3*^*YFP-Cre*^; *R26* ^*CAG-FlpoER/CAG-FlpoER*^ for males and *Brg1*^*F/ΔR*^; *FoxP3*^*YFP-Cre/YFP-Cre*^; *R26* ^*CAG-FlpoER/CAG-FlpoER*^ for females. The rKO mice were fed with a low dose of TAM (12 ug/g, once only, termed the “low dose” regimen hereafter) to reverse *Brg1* KO in a fraction of Tregs. The GFP^+^ and GFP^-^ Treg subsets were then isolated by FACS. As expected, the gene trap cassette was lost in the GFP^-^ subset (Fig. 1D) concomitant with the emergence of the functional *Brg1* transcript (which contained E14-15; Fig. 1E). These data validated the functionality of the Δ*R* allele.

### Dramatic effects of *Brg1* reexpression on rKO mice

The severity of the inflammatory phenotypes was somewhat variable in different rKO mice, and tended to correlate with the frequencies of effector/memory-like (E/M) CD44^hi^CD62L^lo^ CD4 cells in the peripheral blood. For convenience, we used the frequencies of the E/M CD4 cell at 3 weeks of age to divide the rKO mice into two groups: rKO1 (>65%) and rKO2 (< 65%), whose phenotypes are described in Fig. 2A-E and Fig. 2F-H, respectively. Of note, the majority (85%) of the rKO mice belonged to the rKO1 category.

**Fig. 2.**
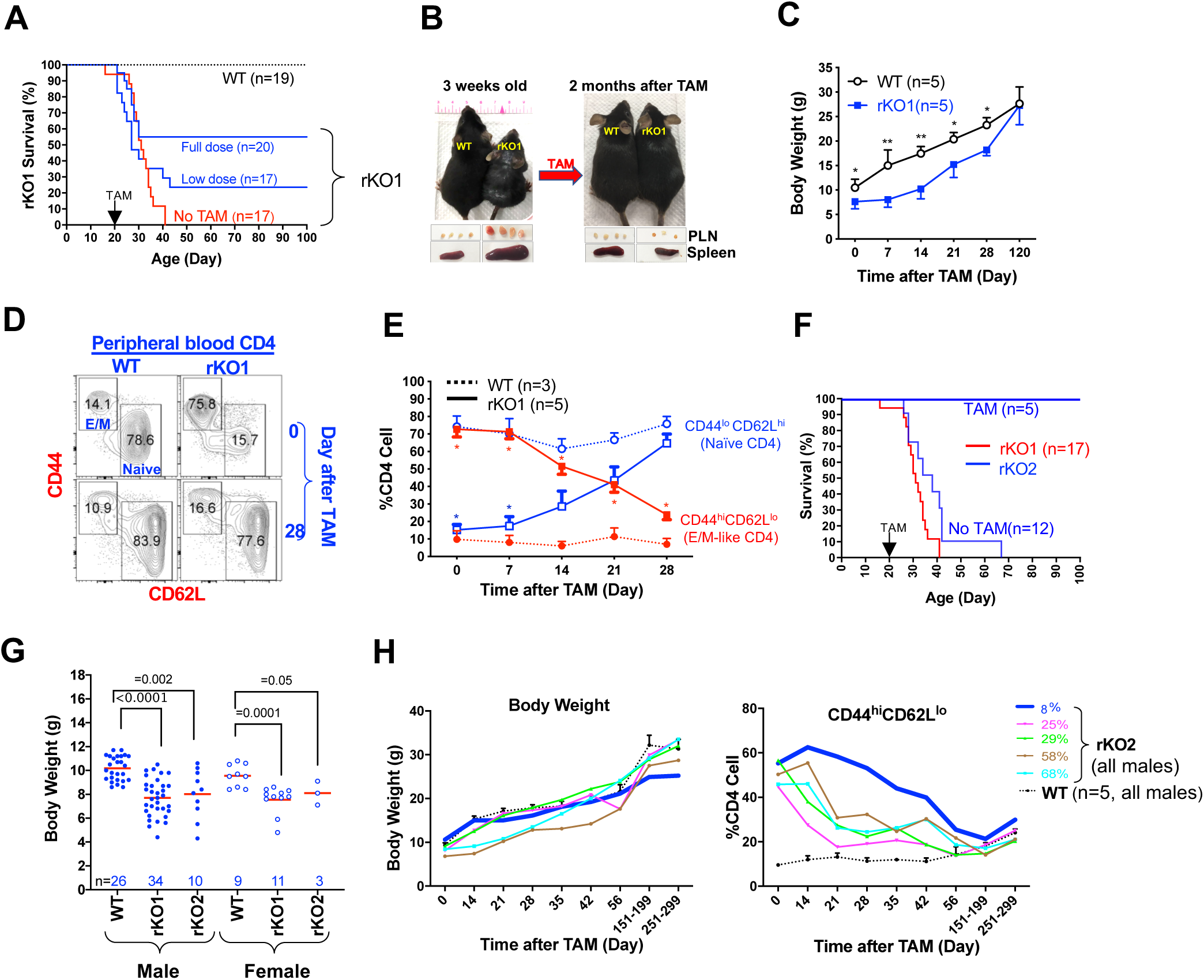
Effects of *Brg1* reexpression on rKO mice. **(A)** Survival of rKO1 mice. rKO1 mice were either *Brg1*^*F/ΔR*^; *FoxP3*^*YFP-Cre*^; *R26*^*CAG-FlpoER/CAG-FlpoER*^ males or *Brg1*^*F/ΔR*^; *FoxP3*^*YFP-Cre/YFP-Cre*^; *R26*^*CAG-FlpoER/CAG-FlpoER*^ females. Littermate controls were of the same genotypes except they carried either *Brg1*^*F/*+^ *or Brg1*^Δ*R*/+^ and thus heterozygous for *Brg1*. For convenience, these controls were labeled “WT” throughout the paper. The rKO1 mice treated with full dose of TAM lived significantly longer than rKO1 (p<0.05). **(B)** Representative image of rKO1 mice before and after TAM treatment (full dose). **(C)** Body weight gain of rKO1 mice following TAM treatment(full dose).* and **, p<0.05 and 0.01, respectively. **(D-E)** Abundance of effector/memory-like (E/M) and naïve CD4 cells in peripheral blood. TAM (full dose) was given to 3-wks-old rKO mice (Day 0). Blue and red asterisks denote statistical significance (p<0.05) when Naïve and E/M CD4 frequencies, respectively, were compared between rKO1 and WT mice. **(F)** Low-dose TAM regimen fully rescued rKO2 mice (p<0.05). For comparison, the survival curve for the rKO1 mice (Fig. 2A) is also displayed (red line). **(G)** rKO2 were nearly as runted as rKO1. **(H)** Body weight (left) and E/M CD4 cells (right) of the 5 TAM-treated rKO2 mice in Fig. 2F. The mean values (+/-SE) of WT littermates are also plotted (dotted lines). As the rKO mice (bearing 5-6 alleles) were quite rare, the 5 mice were born and hence analyzed at different times, except for the last two time points when the mice from different litters were analyzed together (the mouse with 8% reversal analyzed on Day 151 and 251).

We found that the rKO1 mice had developed severe inflammatory signs (including skin lesions, lymphoid organ enlargement and runting) by 3 weeks of age and died before Day 41, the median survival being 31 days (Fig. 2A, red line). To determine the consequences of *Brg1* reexpression in rKO1 mice, mice were given TAM (full dose) around 3 weeks of age, namely ∼10 days before the predicted median death date. Remarkably, 55% of the mice (11/20) were rescued from death (Fig. 2A). Gross signs of inflammation disappeared within two months after TAM administration (Fig. 2B), and by 120 days, the runted mice had fully caught up in weight and size, revealing striking resilience of the mice (Fig. 2C). To directly examine the kinetics of inflammation resolution, we monitored the proportion of effector/memory-like (E/M, CD44^hi^CD62L^lo^) and naïve-like (Naïve, CD44^lo^CD62L^hi^) CD4 cells within the CD4 cell population in peripheral blood (Fig. 2D). In a 3-wks-old rKO1 mouse, the E/M and Naïve subset constituted 76% and 16% of total CD4 population, respectively (as opposed to 14% and 79% in the WT mice; Fig. 2D, top). TAM treatment (full dose) led to pronounced and progressive depletion of the E/M CD4 cells and simultaneous accumulation of the naïve CD4 cells, which became apparent within 2 weeks after the treatment (Fig. 2D-E). The reciprocal changes in the abundance of the E/M vs. naïve CD4 cells were not due to the conversion of the E/M to naïve CD4 cells (Fig. S1), and so might instead reflect the changes in their apoptosis/proliferation rates.

We conclude that reversing *Brg1* KO in all of the *Brg1*-deficient Tregs as late as 10 days before the predicted median death date rescued 55% of the dying mice. However, in clinical settings, it is unfeasible to repair genetic defects in all of the target cells. Therefore, we repeated with the rescue experiment using the low-dose TAM regimen, which resulted in *Brg1*-reexpression in variable fractions (10%-50%) of Tregs among different individuals (not shown). Under this condition, 18% (3/17) of the dying rKO1 mice were rescued (Fig. 2A, low dose), with their inflammation resolved and body weight (largely) recovered (Fig. S2A).

*Brg1*-reexpression proved more effective in rescuing the rKO2 mice, where inflammation was somewhat less devastating. In the absence of TAM, all but one (11/12) rKO2 mice died before Day 42 and the remaining mouse died on Day 67, with the median survival being 38 days, which was only mildly longer than that rKO1 mice (Fig. 2F). Furthermore, the rKO2 mice were nearly as runted as rKO1 (Fig. 2G). Thus, rKO2 mice were also very sick. Nevertheless, following the low-dose TAM treatment in 3-wks-old mice, which restored *Brg1* expression in 8% to 68% Tregs (measured on Day 14 after the treatment; Fig. 4A, right plot), 100% (5/5) of the rKO2 mice survived (Fig. 2F, blue line), with their body weights catching up and inflammation subsiding over time (Fig. 2H). Remarkably, these changes were observed even in the mouse where *Brg1* expression was restored in only 8% of the Tregs, despite quite severe inflammation before TAM treatment (Fig. 2H, thick blue line). Note that his body weight might never fully catch up, remaining slightly lower than an age- and sex-matched littermate control even on Day 251 after TAM (25.2 vs. 28.8g). Nevertheless, by Day 251, this mouse seemed to have become otherwise perfectly healthy, devoid of any overt sign of illness such as skin lesions and lethargy (not shown).

Collectively, these data reveal powerful effects of *Brg1* reexpression on the sick mice, with as little as 8% of *Brg1*-reexpressed Tregs sufficient for the rescue in some cases.

### *Brg1* reexpression, presumably in conjunction with excessive cytokine stimulation, produced hyperactivated, highly suppressive “SuperTregs”

To characterize *Brg1*-reexpressed Tregs, we treated 3-wks-old rKO1 mice with the low dose of TAM and compared gene expression patterns in *Brg1*-deleted (GFP^+^) vs. *Brg1*-reexpressed (GFP^-^) Treg subsets isolated 7 days after TAM, when the two populations were cleanly distinguishable (Fig. 1F). This analysis would reveal the role of *Brg1* in partially activated Tregs exposed to inflammation. As a control, we addressed the role of *Brg1* in Tregs under the physiological condition. To this end, we compared *Brg1*-deleted (YFP^+^) and *Brg1*-sufficient (YFP^-^) Tregs from the healthy, mosaic females (*Brg1*^*F/ΔR*^; *FoxP3*^*YFP- Cre/+*^; *R26* ^*CAG-FlpoER/CAG-FlpoER*^, where YFP-Cre was expressed in only half of the Tregs due to random X-inactivation; these mice also carried *R26* ^*CAG-FlpoER/CAG-FlpoER*^ just as the rKO1 mice in order to control for any potential nonspecific confounding effects of FlpoER expression when comparing differentially expressed genes between the two strains). As additional controls, we used Tregs isolated from WT mice and from rKO1 mice not treated with TAM, the former being *Brg1*-sufficient while the latter *Brg1*-deficient, therefore comparable to *Brg1*-sufficient Tregs from the mosaic females and the *Brg1*-deficient Tregs from TAM-treated rKO1 mice, respectively. All the mice were 3-4 weeks old when sacrificed.

*Brg1*-deletion in the mosaic females and *Brg1*-reexpression in rKO1 mice on Day 7 after TAM treatment affected 618 and 1352 genes, respectively, with only 241 genes shared, suggesting divergent roles of *Brg1* under the physiological vs. inflammatory conditions (Fig. 3A; see Supplemental Data for complete list of these genes; raw data already deposited). *Brg1*-target genes are of diverse functions, a conspicuous group being related to Treg function (Fig. 3B). These genes can be divided into two categories: the “naïve genes” that are predominantly expressed in naïve Tregs (*Bach2* and *Ccr7*) ^8,19^, and “activation/effector function genes” preferentially expressed in activated/effector Tregs, including *Icos* ^8^, *Tigit* ^20^,*Cxcr3*^21^, *Klrg1*^22^, *Prdm1*^23^ and *Gzmb*^24^. In the *Brg1* KO Tregs within the mosaic females, the “naïve genes” were upregulated, while most of the “activation/effector function genes” repressed, relative to the *Brg1*-sufficient Tregs in both the mosaic females and the WT mice (lane 3 vs. 1-2), confirming that the direct effect of *Brg1* deletion was to inhibit Treg activation ^17^. Interestingly, in the rKO1 mice with severe inflammation, the *Brg1* KO Tregs were partially/weakly activated, with the “naïve genes” repressed and some of the “activation/effector function genes” (i.e.,*Cxcr3, Gzma, Gzmb,Gzmf*) upregulated relative to the *Brg1*-sufficient controls (lane 4 vs. 1-2). These data reinforce the notion that *Brg1* KO impairs Treg activation, which triggers inflammation, leading to a secondary partial/weak Treg activation ^17^. As expected, in the rKO1 mice, following the low-dose TAM treatment which restored *Brg1* expression in a subset of Tregs, the *Brg1*-deficient subset remained mostly unaffected, with the expression pattern comparable to that in the rKO1 mice without TAM treatment (lane 5 vs. 4). In sharp contrast, the *Brg1*-reexpressed Treg subset in these mice became dramatically activated, as revealed by 5-10x repression of naïve genes and 2-14x upregulation of all the activation/effector function genes relative to the partially activated, *Brg1*-deleted subset (lane 6 vs. 5). Thus, *Brg1* reexpression in the rKO1 mice led to Treg super-activation, the resultant super-activated Tregs (“SuperTregs”) presumably highly suppressive. The data also demonstrate that although the *Brg1* target genes were in general highly divergent in the mosaic (healthy) vs. rKO1(inflamed) mice (Fig. 3A), the *Brg1*-controlled transcription program underlying Treg activation was conserved between the two distinct conditions, but with a twist: in the rKO1 mice, BRG1 was able to upregulate the activation markers to much higher levels than in mosaic mice (lane 6 vs. 2), which seemed to reflect (in part) a synthetic effect of cytokine stimulation in the rKO1 mice (see further).

**Fig. 3.**
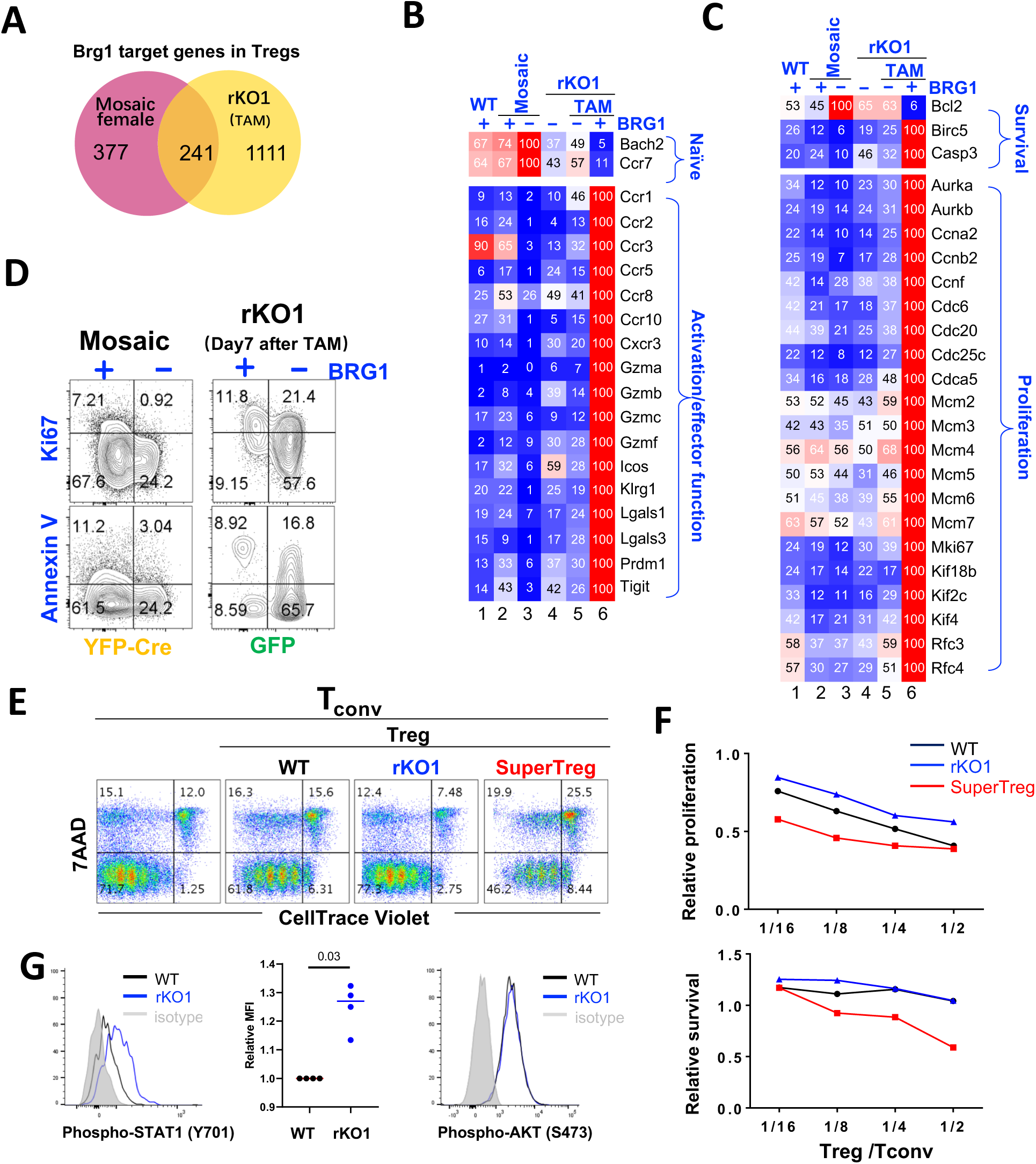
Brg1 reexpression converted dysfunctional Tregs into SuperTregs. **(A)** Brg1-target genes from mosaic females and from rKO1 mice on Day 7 after TAM. The target genes were defined as those whose expression was affected ≥2x fold by BRG1. **(B-C)** Relative expression of representative Brg1 target genes in SuperTregs (lane 5-6), compared with their expression in other Treg types (lane 1-4). These genes were associated with Treg function (B) or turn-over (C). **(D)** FACS analysis of proliferation and apoptosis in the Tregs from mosaic females (left) and from the rKO1 mice on Day 7 after TAM (right). The two mice were analyzed at different times. **(E-F)** Treg suppression in vitro. CD4 cells, labeled with CellTracer, were stimulated with APC and anti-CD3 in the presence or absence of Tregs for 5 days before analyzing 7AAD and CellTracer fluorescence (E). The CellTracer MFI of the viable (7AAD^-^) cells in the presence of Tregs were plotted relative to that in the absence of Tregs, as are the relative viable cell frequencies plotted (F). In E, the Treg: Tconv ratio was 1:4. A replica of this experiment is shown in Fig. S3. **(G)** IFNg -STAT1 and TCR-AKT signaling in splenic Tregs. To quantify STAT1 (Y701) phosphorylation, the STAT1 (Y701) MFI for rKO1 Tregs was normalized to the WT controls, the latter set as 1 (middle).

Activated Tregs are more apoptotic and proliferative than naïve Tregs ^8^. Indeed, in SuperTregs, the prosurvival gene *Bcl2* was repressed whereas the pro-apoptosis gene *Casp3* and many cell cycle promoting genes upregulated when compared with all other Treg types examined (Fig. 3C, lane 6 vs 1-5). Consistent with the increased proliferation, *Brg1*-reexpressed Tregs became somewhat more abundant after TAM treatment (not shown). Curiously, the pro-survival *Birc5* was also upregulated in SuperTregs, perhaps reflecting a negative feedback effect (Fig. 3C).

We next used FACS to validate the RNA-seq results. SuperTregs were indeed more proliferative and more apoptotic (Fig. 3D). To assay their suppressive function, dye-labeled conventional CD4 cells (Tconv) were stimulated with antigen presentation cells and anti-CD3 for 5 days in the presence or absence of Tregs before the FACS analysis. In the absence of Tregs, 72% of the Tconv survived and had undergone multiple rounds of division as revealed by progressive dye dilution (Fig. 3E, left FACS plot). Tregs from WT mice, *Brg1-* deleted Tregs from rKO1 mice and SuperTregs all inhibited proliferation and promoted apoptosis in a dose-dependent manner, but importantly, SuperTregs was the most effective (Fig. 3E-F; see Fig. S3 for a replica experiment).

Finally, we have begun to define the mechanism underlying gene hyperactivation in SuperTregs, namely, how BRG1 in the inflamed rKO1 mice could upregulate the activation markers to much higher levels than BRG1 in the healthy mosaic or WT mice (lane 6 vs. 1-2). To address this issue, we used *Cxcr3* as a model. *Cxcr3* marks the Treg subset specialized in suppressing the Th1 response^21^. It is a direct target of *BRG1* ^17^ and hyperactivated in SuperTregs (Fig. 3B). Importantly, *Cxcr3* is induced in response to IFN*γ* stimulation, perhaps in conjunction with TCR signaling^21^. Given that *Cxcr3* is subject to joint regulation by *BRG1* and IFN*γ* /TCR, our hypothesis is that in rKO1 mice with severe inflammation, Tregs experience enhanced IFN*γ* /TCR stimulation, which can conceivably complement BRG1 to induce strong *Cxcr3* expression. Indeed, in ∼3 weeks-old rKO1 mice with severe inflammation, STAT1 phosphorylation in Tregs was markedly elevated, indicating excessive IFN*γ* signaling (Fig. 3G, left and middle). Interestingly, TCR signaling seemed unaltered in these Tregs (Fig. 3G, right).

Our data collectively suggest that *Brg1* reexpression acted (partly) in conjunction with inflammatory cytokines to convert *Brg1*-deleted Tregs into hyperactivated Tregs endowed with potent suppressive activity.

### The fate of SuperTreg *in vivo*

We have followed SuperTregs in the five TAM-treated rKO2 mice (Fig. 2H); these mice, treated with the low-dose TAM regimen, harbored both GFP^-^ and GFP^+^ Treg subsets, the former being SuperTregs while the latter serving as an internal control for FACS analysis. Peripheral blood was drawn and crucial Treg markers (KLRG1, ICOS, TIGIT, CXCR3, all induced on SuperTregs; Fig. 3B) monitored over time.

In the five mice, SuperTregs comprised 8% to 68% of total Tregs in the blood on Day 14 after TAM (Fig. 2H). We were especially intrigued in the mouse harboring the least (8%) amount of SuperTregs (thick blue line, Fig. 2H). In this particular mouse, the KLRG1^+^ Treg subset, barely detectable within *Brg1*-deleted (GFP^+^) Treg subset, accounted for as much as 21% of the SuperTreg population on Day 14 after TAM, which remained elevated thereafter, presumably reflecting the persistence of certain degree of inflammation (Fig. 4A, row 2-4; Fig. 4B, top left, pink line). By Day 14 after TAM, ICOS had been dramatically induced in SuperTregs, being expressed on (almost) all KLRG1^+^and KLRG1^-^ subsets (as opposed to 32% in *Brg1*-deleted Tregs; Fig. 4A, row2, column 4). Of note, in the ICOS^+^ subset of SuperTregs, the level of ICOS expression was also elevated relative to ICOS^+^ Treg subset in the WT mice, suggesting that SuperTregs were more active than the activated Treg subset in WT mice on the single cell basis (Fig. 4A, row 2, column 3 vs. 1). Interestingly, in contrast to KLRG1, ICOS expression in SuperTregs (and in *Brg1*-deleted Tregs) declined over time to the baseline by Day 251 after TAM, occurring faster in the KLRG1^-^ subset (Fig. 4A, row 2, column 3-8; Fig. 4B, bottom left), suggesting (partial) resolution of inflammation. Indeed, TIGIT and CXCR3, also induced on SuperTregs (albeit to less extents than ICOS), had similarly declined to (near) basal levels by Day 251 (Fig. 4A, row 3-4; Fig. 4B, bottom), as were the abundance of the E/M CD4 cells in the peripheral blood (Fig. 2H, right). Of note, on Day 151 after TAM, the frequency of GFP^-^ Treg subset within the Treg population was markedly increased (to 20.6% from 9.5% on Day 56, Fig. 4A, row 1). To determine whether this increase was due to the accumulation of the GFP^-^ Treg subset and/or depletion of the GFP^+^ Treg subset, we examined the abundance of the two Treg subsets relative to that of conventional CD4 cells, finding that the increased frequency of the GFP^-^ subset was due to its accumulation, as the abundance of the GFP^+^ subset remained constant as compare with Day 56 (Fig. 4B, right, heavy lines). Interestingly, by Day 251, GFP^+^ subset had become partially depleted while the GFP^+^ subset further accumulated (Fig. 4B, right, heavy lines, last time point). The mechanisms underlying the reciprocal changes in the two Treg subsets are unclear, but might involve competition between the *Brg1*-sufficient and *Brg1-*deficient Tregs.

**Fig. 4.**
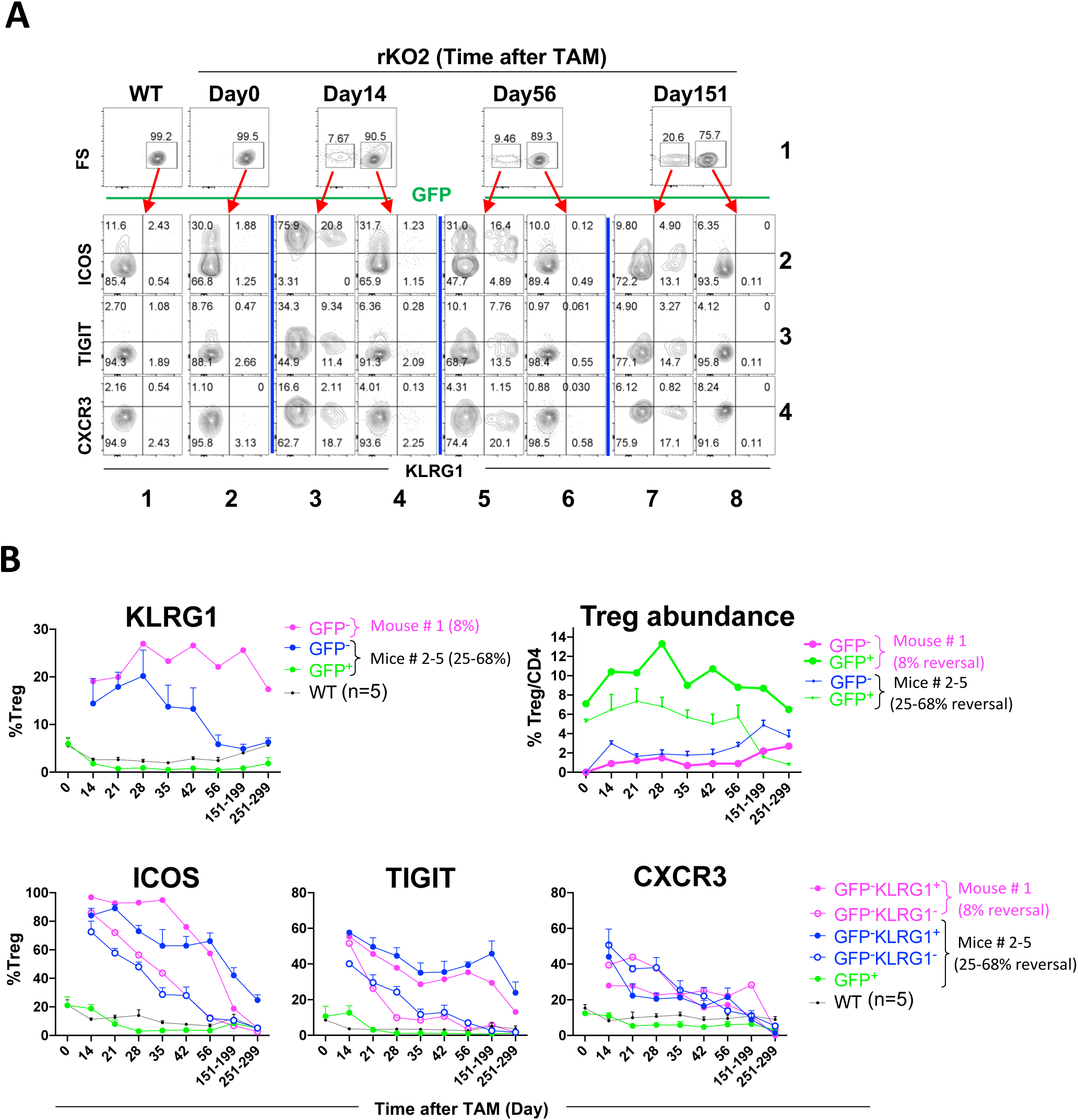
Fate of SuperTregs *in vivo*. **(A)** Peripheral blood cells from the rKO2 mouse with 8% GFP^-^ Tregs (Fig. 2H, thick blue line) were stained with a mixture of CD4, KLRG1, ICOS, TIGIT and CXCR3 antibodies for the analysis of ICOS, TIGIT and CXCR3 expression in KLRG1^+^ and KLRG1^-^ subsets (row 2-4) within the GFP^-^ (column 3,5,7) and GFP^+^ (column 1,2,4,6,8) Treg populations. **(B)** Summary of the FACS results for the mouse analyzed in Fig. 4A (mouse #1), together with the four other mice in Fig. 2H (#2-5). For clarity, only the GFP-subset is displayed for mouse #1 in all the plots except the top right plot, and only Mean +/- SM is displayed for Mice #2-5. As the rKO mice (bearing 5-6 alleles) were quite rare, the 5 mice were born and hence analyzed at different times, except for the last two time points when the mice from different litters were analyzed together (the mouse with 8% reversal analyzed on Day 151 and 251).

In the remaining four rKO2 mice (#2-4), where *Brg1* was reexpressed in more (25% to 68%) Tregs (Fig. 2H), the E/M CD4 cells were depleted far more rapidly (Fig. 2H, bottom), and all the activation markers (including KLRG1) decayed over time (Fig. 4B), consistent with more effective resolution of inflammation. The reciprocal changes in the abundance of the GFP^-^ vs GFP^+^ Treg subsets were also observed (Fig. 4B, top right, thin lines). Finally, we also followed the fate of the 3 rKO1 mice treated with the low-dose TAM regimen, with similar findings (Fig. S2B).

We conclude that SuperTregs tended to lose the hyper-activated phenotype as the inflammation subsided, suggesting that the inflammatory environment was essential for maintaining Treg hyperactivation. Our data also support the notion that the enhancement of Treg suppressive function in response to inflammation is “memory-less”, a feature important for avoiding generalized immunosuppression that could otherwise result from repeated activation^9^.

## DISCUSSION

Using conventional gene KO technologies, many genes have been identified that affect Treg function and immune tolerance. The current work is the first to address the effects of reversing the KO, which provides insights hard to obtain using conventional KO models, as discussed below.

### Consequences of *Brg1* KO and reexpression in Tregs

These are summarized in Fig. 5, which is based on the current and the previous work^17^. Specifically, in WT mice, when antigens activate conventional T cells, Tregs also get activated to restrict the immune response. In rKO mice, *Brg1* KO impairs Treg activation, leading to the onset of inflammation. As the inflammation intensifies, Tregs get partially activated (partly) by inflammatory stimuli such as IFN*γ*, but this is insufficient to stop the ongoing inflammation (dotted line). Importantly, at this point, when *Brg1* is re-expressed upon TAM administration, it acts in conjunction with the inflammatory stimuli to convert the functionally compromised Tregs into hyperactivated “SuperTregs”, which overwhelm the inflammation. As the inflammation is resolved, SuperTregs reverse activation-induced changes (not depicted).

**Fig. 5.**
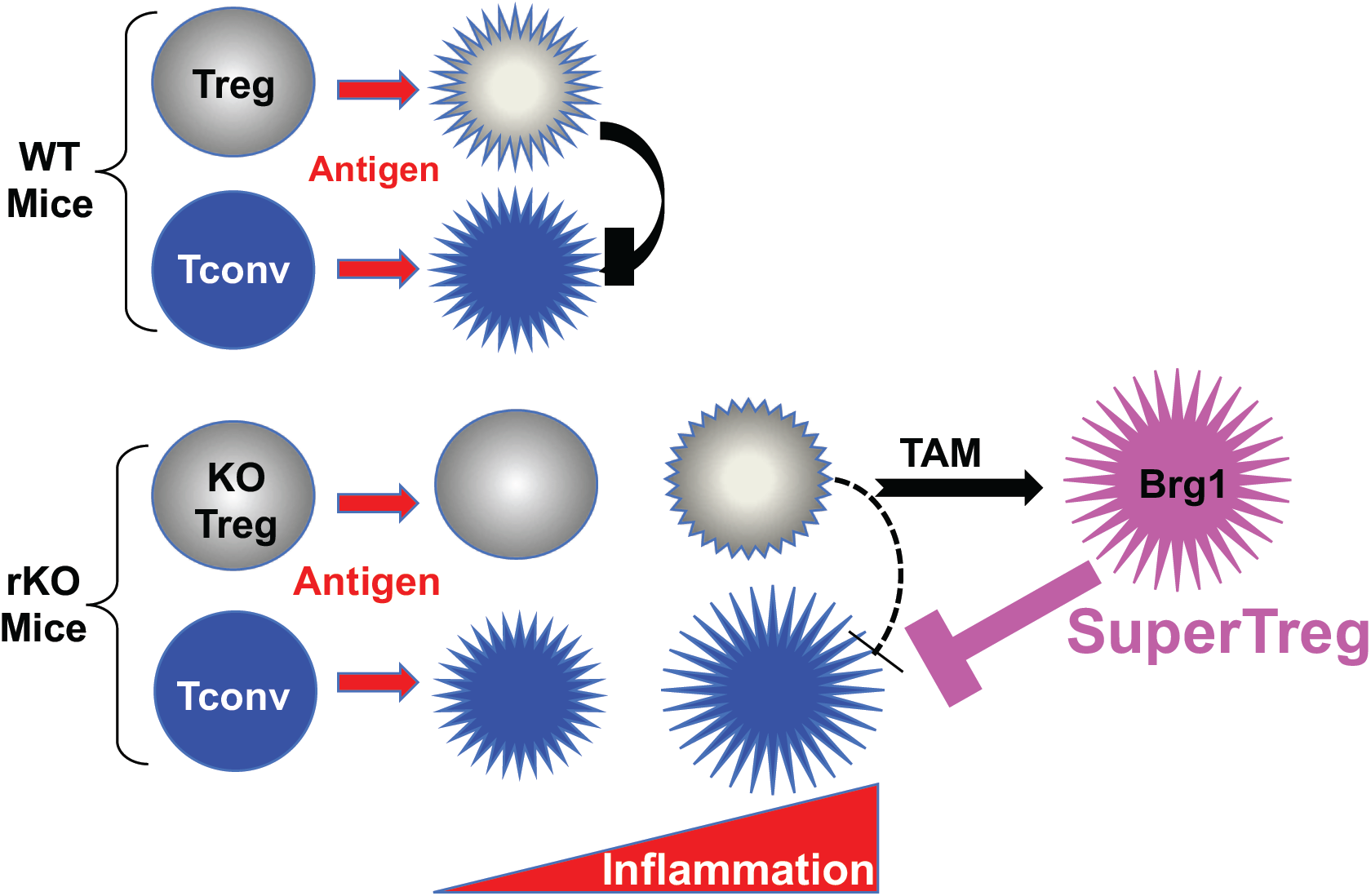
A model. See text for detail.

A few issues are noteworthy regarding this model.

First, we wish to reiterate that compared with the activated Tregs in the WT mice, the activated Tregs in the SuperTreg population were not only more abundant, but also expressed higher levels of some activation markers (like ICOS). Thus, SuperTregs showed both quantitative and qualitative differences from the activated Tregs in the WT mice.

Second, TAM treatment should also lead to *Brg1* reexpression in the Treg precursors in the thymus and bone marrow, and the nascent, *Brg1*-sufficient Tregs might also contribute to the resolution of inflammation. However, this contribution might be minimal, given the low rate of T cell production in adult mice, especially in the sick mice where the thymi were profoundly atrophic as a result of inflammatory stress (not shown).

Third, Brg1-reexpressed Tregs had markedly accumulated by 5 months after the low-dose TAM treatment, which should ensure permanent benefit of the treatment.

Finally, excessive STAT1 signaling (caused e.g. by Treg-specific SOCS1 ablation) is known to cause CXCR3 overexpression, which paradoxically impairs the ability of Tregs to control Th1 response, in apparent conflict with our observation^25^. We note that the CXCR3 is much more overexpressed in the SOCS1-deleted Tregs than in our SuperTregs (Fig. 6H in reference vs. Fig. 4 in this study). Perhaps STAT1 signaling can produce opposite effects when elevated to different levels.

### Value of reversible KO mice

In contrast to our previous study which used conventional gene KO model to address the role of *Brg1* in naïve Tregs in normal mice, the current study used a reversible KO method to explore the consequence of *Brg1* reexpression in the sick mice. Obviously, the effect of *Brg1* reexpression on the partially activated Tregs was hard to predict from the known roles of *Brg1* in naïve Tregs, partly because of the change in the information context of *Brg1* action under the two distinct conditions (see Introduction); indeed, the majority of *Brg1*-affected genes in SuperTregs were different from that affected by *Brg1*-deletion in naïve Tregs (Fig. 3A). It is even harder to predict that *Brg1*-reexpression in as little as 9% of defective Tregs would suffice to resolve inflammation in the sick mice; indeed, to our knowledge, the efficacy of Tregs to stop severe ongoing systemic inflammation in adult/adolescent mice remains largely unexplored, although it is well known that adoptive transfer of Tregs into neonatal *Scurfy* mice (which lack Tregs) can prevent the onset of lethal autoimmunity (see, e.g, ^21^. Our study thus illustrates the value of reversible KO methods in uncovering gene functions. Unfortunately, such methods remain way underutilized, despite brilliant successes in a few isolated cases published in high profile journals^26–28^.

### Medical relevance of the current study

Our study has therapeutic implications for heritable autoimmune disorders resulting from Treg defects, the best defined being the immune dysregulation, polyendocrinopathy, enteropathy, X-linked (IPEX)resulting from *FOXP3* mutations^3–5^. The IPEX phenotypes tend to vary with the nature of the mutations. For example, missense mutations and promoter mutations can be associated with normal Treg numbers (but compromised Treg suppressive function) and a milder phenotype. In addition to FoxP3, mutations at a number of other genes important for Treg function (including *CD25, STAT5b, ITCH* and *STAT1*) are known to cause IPEX-like disorders ^5^. Treatment options for the IPEX disorder are limited mainly to immunosuppressive drugs and allogeneic hematopoietic stem cell transplantation (HSCT). Immunosuppressive therapy is beneficial only temporarily, as it fails to prevent disease progression in most patients, with the overall survival rate being only 65% at 24 years of age^4^. HSCT does not improve the survival rate, and furthermore, some patients cannot undergo HSCT due to limited donor availability or because their clinical manifestations are not severe enough to justify HSCT ^4,29^. Effective therapies for IPEX-like disorders similarly remain elusive.

We envision an alternative strategy for treating IPEX(-related) disorders. In contrast to HSCT, our strategy exploits preexisting defective Tregs. Specifically, we propose to correct the genetic defects in the Tregs *in vivo*, thus restoring their function and even converting them into SuperTregs. This conversion is plausible if the mutations compromise Treg activation in a reversible manner as in the case of *Brg1* KO. Alternatively, the mutations might not affect Treg activation but block some other aspects of Treg function. In this case, the defective Tregs should already be activated in the inflammatory environment prior to gene therapy, and if the particular Treg defects are (partially) reversible, then repairing the mutations might suffice to convert the Tregs into SuperTregs, which seems feasible at least for the IPEX patients with normal numbers of Tregs mentioned above.

The efficiency of gene-editing determines the therapeutic efficacy. Gene editing tools vary in efficiency. Fortunately, the highly effective “base editors” that can change A>G or C>T have been developed ^30,31^, which is applicable to, for example, the many IPEX patients carrying a single G>A substitution at FoxP3^4^. The base editor together with relevant gRNA expression cassette might be delivered systemically into such patients using a lentiviral vector, such as the CD4-targeted lentiviral vector that transduces up to 7% of human CD4 cells in mice following a single i.p injection ^32,33^. This strategy may particularly benefit patients with the Tregs mildly compromised in function but normal in numbers, where correction of the mutations in a small fraction of these Tregs might suffice to effect a cure. This gene-editing based strategy may not be far-fetched. Indeed, In animal models, gene editing has shown great promises for treating monogenic diseases (via simple injection of gene editing components) such as Duchenne Muscular Dystrophy and Leber Congenital Amaurosis 10, the latter already approved for Phase ½ trials^34,35^. IPEX(-related) disorders represent valid candidates for gene editing-based therapies, as previously proposed ^3^.

## MATERIALS AND METHODS

### Mice

*Brg1*^*ΔR*^ allele was generated using traditional gene targeting strategy as described for the *Baf57*^*ΔR*^ allele^18^, except that the homology arms in the *Baf57*^*ΔR*^ targeting construct were replaced with the sequences from the *Brg1* locus (Fig. 1A). The rKO mice were then created by introducing *Brg*^*F* 36^, *R26*^*CAG-FlpoER* 37^ and *FoxP3*^*YFP-Cre* 38^ into the *Brg1*^Δ*R*/+^ mice. Of note, this breeding scheme also generated conventional, irreversible KO littermates, whose genotypes were identical to rKO except that both alleles of *Brg1* were floxed. Interestingly, the phenotype of these littermates were generally weaker than rKO mice (but similar to the conventional *Brg1* KO mice previously described^17^, presumably because the conventional KO mice carried two copies of *Brg*^*F*^, both of which must be deleted to eliminate *BRG1*, whereas in rKO mice, Cre only needed to delete a single *Brg*^*F*^. The mice were maintained on C57/B6 background. All the experiments were approved by the animal ethical committees at ShanghaiTech and Yale University, and were performed in accordance with institutional guidelines.

### Tamoxifen (TAM) treatment

For full dose regimen, 50 mg TAM (Sigma Aldrich) was added to 900ul corn oil plus 100ul 100% ethanol (50 mg/ml final concentration), and dissolved by incubation at 55°C for 30 min. The solution can be stored at −20°C. The drug was delivered (typically into 3-wks-old mice) via oral gavage at 10 ul/g body weight, once a day for two consecutive days. Low-dose regimen was identical except that the drug was at lower concentration (1.25 mg/ml) and delivered by a single gavage, translating to 40x less TAM as compared with the full dose regimen. While the full dose regimen invariably caused complete deletion of the gene-trap cassette, the low-dose regimen produced variable, highly unpredictable deletion, with the efficiencies ranging from 7% to 70% in difference individuals.

### Flow cytometry

Lymphocytes were stained with antibodies and analyzed using FACS fortessa (BD Biosciences). Phospho-STAT1 and phospho-AKT in splenic Tregs were detected using the following cocktail and the Transcription Factor Buffer Set (BD Pharmingen, 562574):CD4-BV650 or CD4-APC (Biolegend), FoxP3-Percp5.5(BD), Stat1 (pY701)-PE-Texas Red (BD) and AKT (pS473)-BV421 (BD). To minimize sample-to-sample variation of Phospho-STAT1 and phospho-AKT signals, WT and rKO1 splenocytes were stained with CD4-BV650 and CD4-APC respectively before the cells were pooled and stained with the remaining antibodies. The cells in Fig. 4 were analyzed with the following cocktail: CD4-BV650 (Biolegend), CD25-BV605(Biolegend), CD278-PE(Biolegend), CXCR3-APC (eBiosciences), KLRG1-BV421 (BD), TIGIT-APC-R700 (BD).

### Gene expression profiling by RNA-seq

Lymphocytes from lymph nodes and spleens from 3-4-wks-old mice were first magnetically depleted of non-CD4 cells before electronic sorting of Tregs (CD4^+^CD25^+^YFP^+^). Total RNA was isolated from 0.1 million Tregs using RNAprep Pure Micro Kit (TIANGEN), and cDNA synthesized from mRNA using SMART-Seq® v4 Ultra™ Low Input RNA Kit (Clontech). Library was then constructed and sequenced on Illumina HiSeq platform with the PE150 strategy, which yielded 25∼ 60 million reads per sample. To identify differentially expressed (DE) genes between the *Brg1*^+^ and *Brg1*^*-*^ Treg subsets in mosaic mice and TAM-treated rKO1 mice, the count data were TMM normalized, the genes < 5 cpm for both subsets filtered out, and the *p* values adjusted by the Benjamini and Hochbergare method. DE genes are defined as those with absolute fold-changes ≥2 and *padj*<0.05. The data have been deposited (BioProject ID PRJNA547476):https://dataview.ncbi.nlm.nih.gov/object/PRJNA547476?reviewer=r0talgg7e1c3r0nmh68gm2r8bj

### *In vitro* suppression assay

Conventional CD4 cells and Tregs were isolated from PLN and spleens from 3 to 4-wks-old mice. CD4^+^ cells were first enriched using Mouse CD4 T Cell Isolation Kit (Biolegend) before electronic sorting. *Brg1* KO Tregs and SuperTregs were isolated from rKO1 mice before and 7 days after TAM, respectively, while *Brg1*-sufficient littermates (*Brg1*^*F/+*^; *FoxP3*^*YFP-Cre(/YFP-Cre)*^; *R26*^*CAG-FlpoER/CAG-FlpoER*^) used as the source of conventional CD4 cells (CD4^+^CD25^-^ YFP^-^) and WT Tregs (CD4^+^CD25^+^YFP^+^). The purity of conventional CD4 and Tregs exceeded 95% and 90%, respectively. To assess Treg function, conventional CD4 cells (5×10^4^) were labeled with CellTrace Violet (GIBCO) and stimulated with *Rag1*^*-/-*^ splenocytes(5×10^4^) plus 1ug/ml anti-CD3e in the presence of indicated numbers of Tregs. Five days later, the cells were stained with 7^-^AAD and anti-CD4 APC before flow cytometrical analysis of proliferation and survival of the conventional CD4 cells.

## STUDY APPROVAL

All mouse studies were approved by the IACUC at the Shanghai Institute of Biochemistry and Cell Biology, Chinese Academy of Sciences, and conducted in an AAALAC-accredited facility in compliance with the relevant regulations.

## CONFLICTS OF INTEREST

The authors declare no conflict of interest.

## ACKNOWLEDGEMENT

We thank T. Chatila, L. Lu, D. Rudra and Y. Wan for advice.

---

**Fig. S1.**
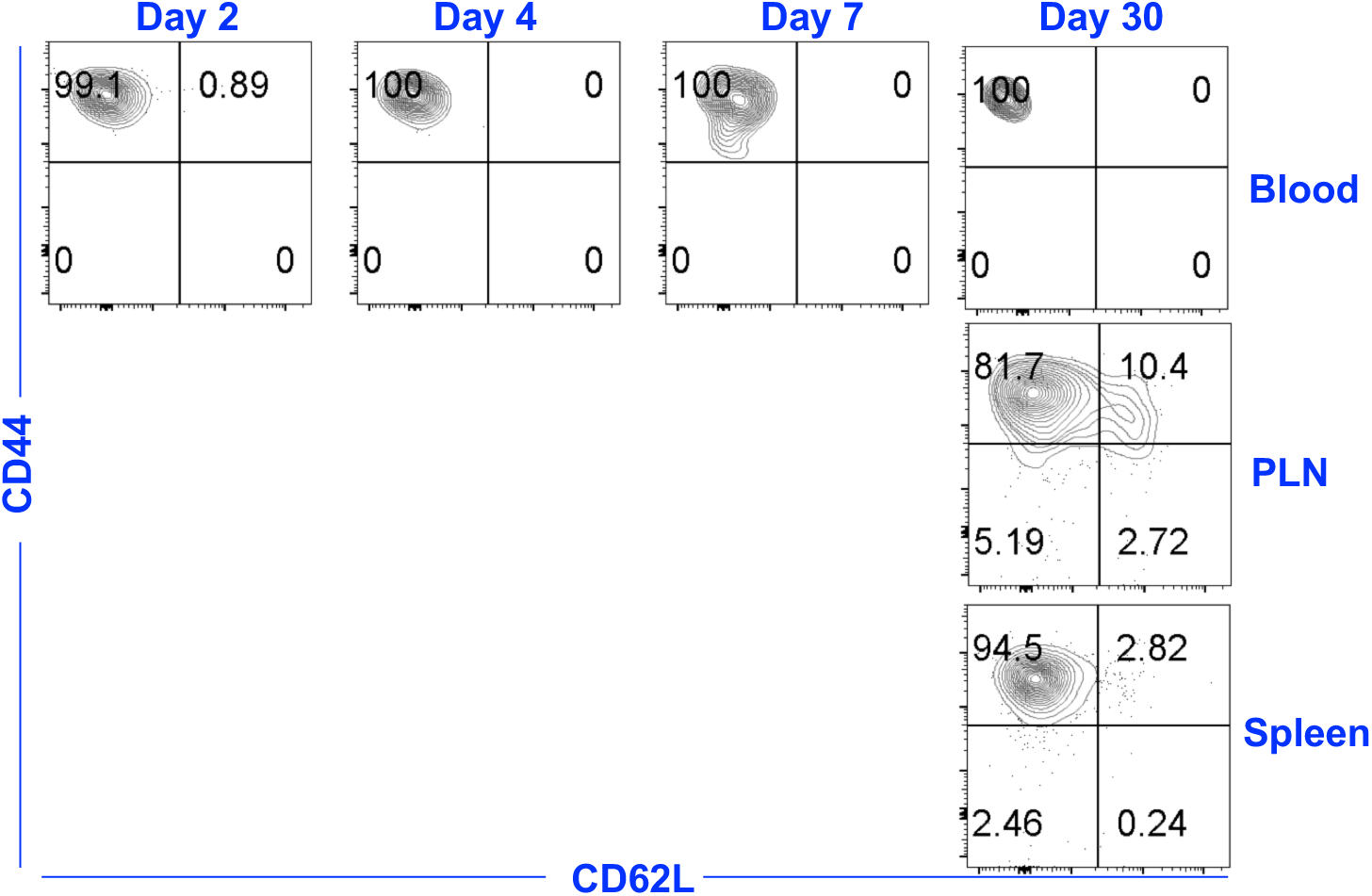
E/M CD4 did not change to naïve CD4 *in vivo*. E/M CD4 cells were isolated from rKO1 mice, labeled with Celltracer Violet before adoptive transfer into an rKO1 mouse just treated with the full dose of TAM. The labeled E/M cells remained CD44^hi^CD62L^lo^ in the blood, and some of the them in the LN and spleen expressed CD62L one month after the transfer. However, none of them was converted to CD44^lo^CD62L^hi^ naïve CD4 cells.

**Fig. S2.**
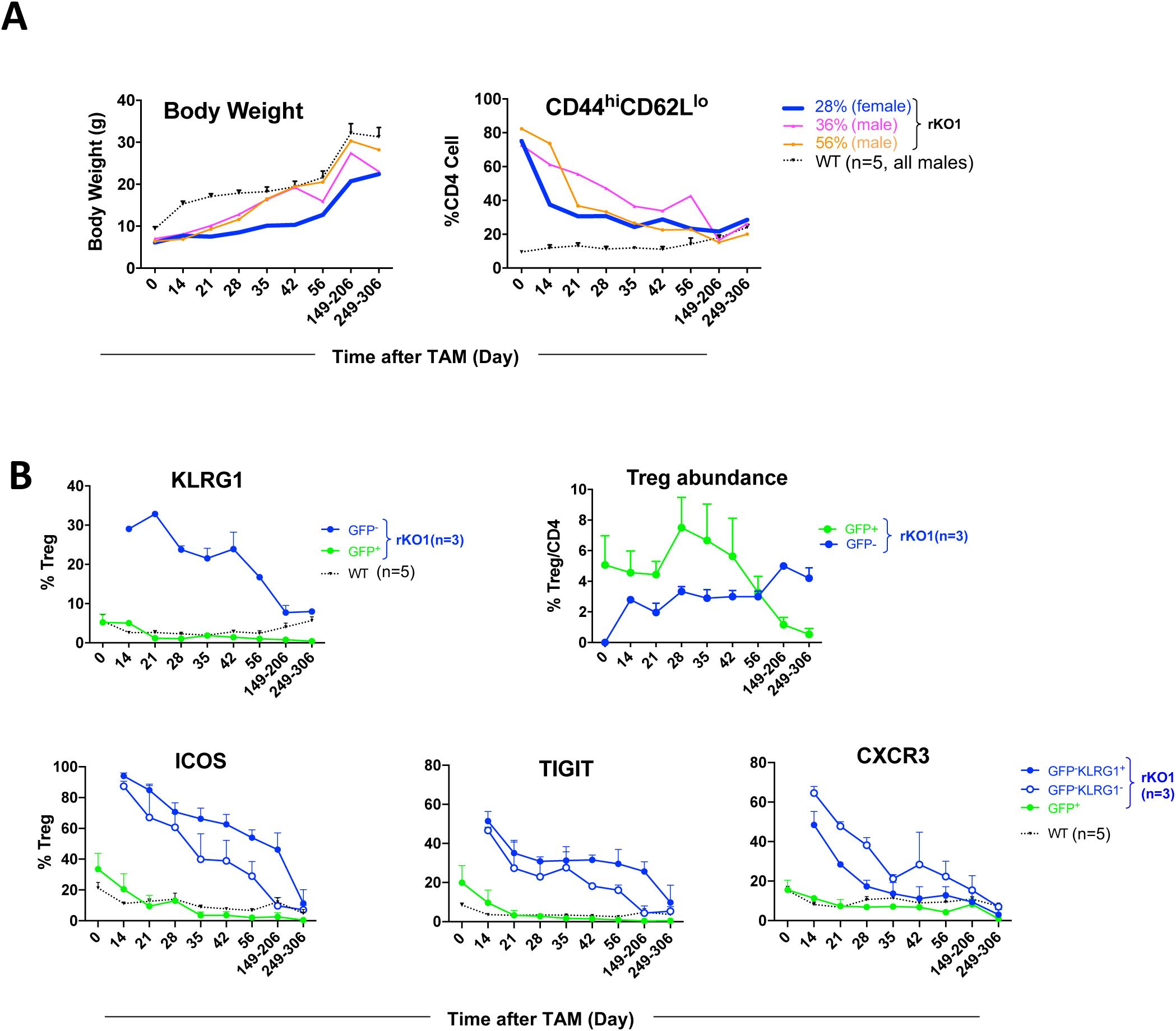
Analysis of the three rKO1 mice rescued by TAM (low dose) **(A)** Body weight and blood E/M CD4 cells, analyzed as for rKO2 in Fig. 2H. The mouse with least *Brg1* re-expressed Tregs (28%) is highlighted with thick blue line. As the rKO mice (bearing 5-6 alleles) were quite rare, the 3 mice were born and hence analyzed at different times, except for the last two time points when the mice from different litters were analyzed together. The control mice were the same as those used for the rKO2 mice (Fig. 2H; Fig. 4). (B) SuperTreg fate, analyzed as for rKO2 in Fig. 4.

**Fig. S3.**
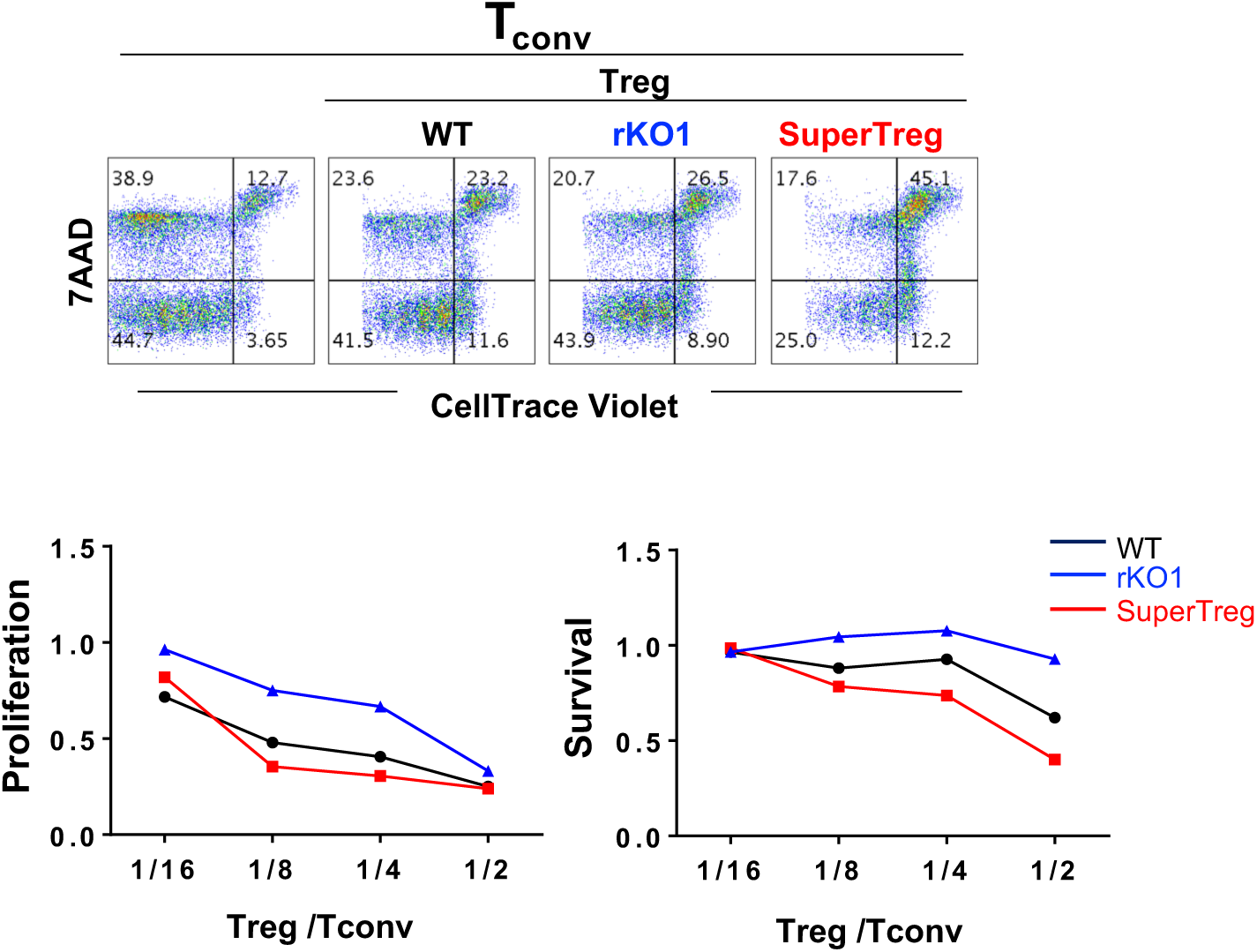
*In vitro* suppression assay, which is a repetition of the experiment in Fig. 3E-F.

